# Multiparametric sensing of outer membrane vesicle-derived supported lipid bilayers demonstrates the specificity of bacteriophage interactions

**DOI:** 10.1101/2022.12.13.520201

**Authors:** Karan Bali, Zixuan Lu, Reece McCoy, Jeremy Treiber, Achilleas Savva, Clemens F. Kaminski, George Salmond, Alberto Salleo, Ioanna Mela, Rita Monson, Róisín M. Owens

**Affiliations:** Department of Chemical Engineering and Biotechnology, University of Cambridge, CB3 0AS Cambridge, United Kingdom; Department of Materials Science and Engineering, Stanford University, Stanford, CA, USA; Department of Biochemistry, Hopkins Building, Downing Site, Tennis Court Road, Cambridge, CB2 1QW, United Kingdom

**Author notes:** **Corresponding Author**, Professor Róisín M. Owens.

**Keywords:** Outer membrane vesicles, supported lipid bilayer, fluorescent microscopy, QCM-D, PEDOT:PSS, bacteriophage-membrane interactions

## Abstract

The use of bacteriophage, viruses that specifically infect bacteria, as antibiotics has become an area of great interest in recent years as the effectiveness of conventional antibiotics recedes. The detection of phage interactions with specific bacteria in a rapid and quantitative way is key for identifying phage of interest for novel antimicrobials. Outer membrane vesicles (OMVs) derived from gram-negative bacteria can be used to make supported lipid bilayers (SLBs) and therefore *in vitro* membrane models that contain naturally occurring components of the bacterial outer membrane. In this study, we used *Escherichia coli* OMV derived SLBs and use both fluorescent imaging and surface sensitive techniques to show their interactions with T4 phage. We also integrate these bilayers with microelectrode arrays (MEAs) functionalised with the conducting polymer PEDOT:PSS and show that the pore forming interactions of the phage with the SLBs can be monitored using electrical impedance spectroscopy. To highlight our ability to detect specific phage interactions, we also generate SLBs using OMVs derived from *Citrobacter rodentium*, which is resistant to T4 phage infection, and identify their lack of interaction with phage. The work presented here shows how interactions occurring between phage and these complex SLB systems can be monitored using a range of experimental techniques. We believe this approach can be used to identify phage against bacterial strains of interest, as well as more generally to monitor any pore forming structure (such as defensins) interacting with bacterial outer membranes, and thus aid in the development of next generation antimicrobials.

---

It is estimated that 25 000 people in Europe die every year from antibiotic resistant bacteria^1^. This number is rising and it is projected that antibiotic resistance could be the cause of 10 million global deaths by 2050^2^. The problem is compounded by the slowdown in effective development of antibiotic treatments. In the last 30 years, there has been a 90% reduction in novel antibiotics approved by the US Food and Drug Administration (FDA)^3^. Out of this concerning landscape, dominated by the dual issues of increasing antibiotic resistance and decreasing novel antibiotic development, bacteriophage therapy as arisen as an alternative strategy. Bacteriophage, or phage for short, are viruses that specifically infect bacteria. They are the most abundant biological entities on earth, with an estimated 4.8 × 10^31^ phage particles in the whole biosphere^4^. Phage are capable of infecting a specific range of bacterial strains depending on the components existing in the cell membrane of these bacteria. This specificity, coupled with the fact phage cannot infect eukaryotic cells means that phage therapy can be a highly precise form of antibiotic treatment with minimal side effects^5^.

A hurdle to the usage of phage therapy becoming more prevalent is the screening of interactions between phage and bacteria. Currently, the main method used to identify and enumerate phage interactions with specific bacteria is the double agar overlay assay^6^. This assay relies on mixing phage and bacterial cultures together with soft agar; if the phage can infect the bacteria, clear spots (plaques) appear on the bacterial lawn. Although this technique has been used for many years, it is time consuming and not suited to rapid phage screening. Therefore, new methods are required that seek to provide a simpler way to precisely evaluate any given phage-host interaction.

The ability of a phage particle to infect a bacterial cell relies first and foremost on the interaction made with specific components in the outermost layer of the cell^7^. In the case of Gram-negative bacteria, whose members make up the bulk of the antimicrobial resistant strains, this outermost layer is the outer membrane (OM)^8^. The OM consists of an asymmetric lipid bilayer with phospholipids in the inner leaflet and glycolipids, predominantly lipopolysaccharides (LPS), in the outer leaflet^9^. The OM also contains outer membrane proteins (OMPs) which play a variety of roles in cellular functioning, notably in the permeation of specific molecules^10^. Phage have tail structures which recognize components of the LPS, OMPs or both^11^, attaching themselves to the cell and initiating membrane penetration and genome injection. For instance, the well-characterised T4 phage binds to both OmpC and LPS on the surface of *Escherichia coli* prior to infection, whilst T7 phage binds to only LPS molecules^7,12^.

Supported lipid bilayers (SLBs) have been used extensively as tools to investigate membrane interactions in an *in vitro* setting. The strength of the SLB platform lies in its ability to be integrated with a range of measurement techniques to probe the events occurring in these membrane mimic systems. However, SLBs generated from synthetic lipids do not faithfully represent the true structure of a cell membrane and this has been a major weakness. By using outer membrane vesicles (OMVs), extracellular vesicles of diameter 20-250 nm that are naturally produced by Gram-negative bacteria, it has been possible to generate SLBs that contain components (such as LPS and OMPs) that are found in the OM of a cell^13,14^. These OM SLBs have been characterized using high resolution microscopy methods, with their components elucidated by such techniques as structured illumination microscopy (SIM) and atomic force microscopy (AFM)^15^.

A key part of the attraction of the SLB platform is their ability to be integrated with a range of measurement techniques. One such measurement technique is electrochemical impedance spectroscopy (EIS), used to probe the resistance and capacitance characteristics of the SLB. SLBs can be generated upon microelectrode arrays (MEAs), and moreover these MEAs can be functionalized with biocompatible conducting polymers, such as the polymer poly(3,4-ethylenedioxythiophene) polystyrene sulfonate (PEDOT:PSS) to increase the device sensitivity. PEDOT:PSS, owing to its ion-to-electron mixed conductivity and cushioned mechanical properties, is an ideal material for interfacing with biological materials (such as SLBs), leading to the emergence of organic bioelectronic devices^16,17^. OM SLBs on PEDOT:PSS coated MEAs have been probed by EIS previously; the actions of the antibiotics polymyxin B, bacitracin and meropenem with *Escherichia coli* OM SLBs were monitored using this technique^18^. However, the OM SLB has not yet been investigated in the context of bacteriophage interactions.

In this study we use T4 phage, a phage that specifically infects *E. coli* cells, measuring its interactions with SLBs using optical and electrical techniques. The basis for the study relies on the interaction occurring between T4 phage particles and *E. coli* OM components, as shown schematically in Figure 1a. Briefly, the phage first attaches to the OM via interactions with LPS and OmpC components in the OM, and after this initial attachment the phage forms a pore in the OM through which it injects its DNA contents into the host cell. We start by showing the specificity of the infection with whole cells using *Citrobacter rodentium*, bacteria that are a closely related species to *E. coli* but crucially not susceptible to T4 infection, as a negative control. We generated OM SLBs from both *E. coli* and *C. rodentium* derived OMVs and assessed their interactions with T4 phage using Structured Illumination Microscopy (SIM). We further investigated their interaction using Quartz Crystal Microbalance with Dissipation monitoring (QCM-D), a technique that measures surface mass changes with high sensitivity. Finally, we integrated SLBs with PEDOT:PSS coated MEAs and performed EIS, showing that the specific interaction between the T4 phage and *E. coli* OM SLBs can be detected electrically. By combining the impedance signature with the optical data, we show here the ability of the OMV derived SLB as a quantitative phage screening platform which can play a role in furthering the prominence of phage therapy as a viable alternative to conventional antibiotic treatments.

**Figure 1.**
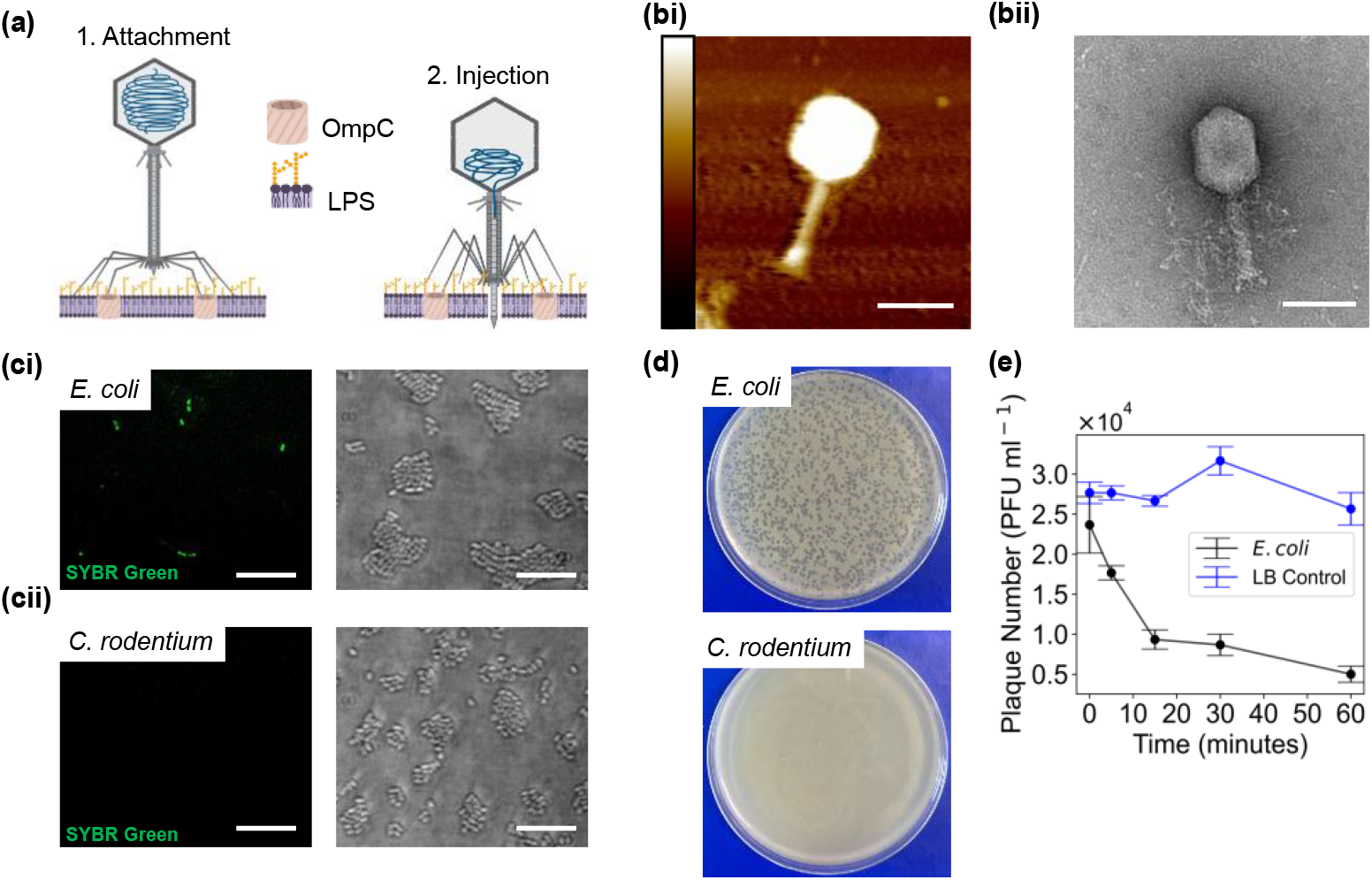
Characterisation of T4 phage whole cell infection. (a) Schematic of the stages of T4 phage interaction with the OM of an *E. coli* cell. (bi) AFM image of a single T4 phage particle. Scale bar = 100 nm; Height bar = 0-36 nm. (bii) TEM image of a single T4 phage particle. Scale bar = 100 nm. (ci) SIM imaging of *E. coli* cells after incubation with SYBR green stained T4 phage, imaged in the (left) 488 nm wavelength and (right) brightfield channels. (cii) SIM imaging of *C. rodentium* cells after incubation with SYBR green stained T4 phage, imaged in the (left) 488 nm wavelength and (right) brightfield channels. (d) Plaque assay in which the same concentration of T4 phage is mixed with *E. coli* and *C. rodentium* cells. The mixture is combined with soft agar and left to set overnight. The plaques seen in the *E. coli* are evidence of T4 infection. (e) T4 phage adsorption assay conducted with *E. coli* cells (m.o.i = 0.01). Samples were taken at regular time intervals; the number of free phage over time were enumerated by spot assay (n = 3) from which the concentration of free phage in PFU ml^−1^ was calculated.

## EXPERIMENTAL

### Bacterial strains and culture conditions

The bacterial strains are shown in the table below:

| Strains | Description | Reference/Source |
| --- | --- | --- |
| *Escherichia coli* BL21 (DE3) | Lab strain | Invitrogen |
| *Citrobacter rodentium* DBS100 | Wild type (WT) strain | GPCS Lab Strain Collection; Popov et al.<sup>19</sup> |

*E. coli* and *C. rodentium* strains were grown at 37 °C in Lysogeny Broth (LB).

### Phage propagation

10 μl of serial dilutions of phage was mixed with 200 μl of overnight *E. coli* culture and 4 ml of molten top agar (0.35% agar), and then poured onto a LB agar plate (1.5% agar) and incubated overnight at 30 °C. Phage concentration was determined by counting plaques on the plate to give the concentration in plaque forming units per ml (PFU ml^−1^). To harvest the phage, the top agar was scraped off the plate, and the surface washed with 3 ml LB. The wash was added to the harvested top agar, and vortexed vigorously with 500 μl chloroform for 2 minutes. The agar mix was then centrifuged at 2220 *g* for 20 minutes at 4 °C. The supernatant was removed, and 10 μl of NaHCO_3_ chloroform was added and vortexed briefly. The phage lysate was stored at 4 °C for further use.

### AFM imaging of T4 phage

AFM images were acquired in Scanasyst mode using ScanasystFluid+ probes (Bruker), with a nominal spring constant of 0.7 N m^−1^ and a resonant frequency of 150 kHz. Images were recorded at scan speeds of 1.5 Hz and tip–sample interaction forces between 200 and 300 pN. T4 phage sample was incubated on mica for 20 minutes before washing and imaging. The resulting images were analysed using Nanoscope to evaluate the T4 phage dimensions.

### SIM imaging of T4 phage

Phage were stained by incubating 1 μl of 1:10 stock concentration SYBR green (Invitrogen) with 200 μl of phage lysate for 15 minutes. Excess dye was removed through a Zeba spin desalting column (Thermo Scientific). To acquire structured illumination microscopy images, a ×60/1.2 NA water immersion lens (UPLSAPO 60XW, Olympus) focused the structured illumination pattern onto the sample, and the same lens was also used to capture the fluorescence emission light before imaging onto an sCMOS camera (C11440, Hamamatsu). The wavelengths used for excitation was 488 nm (iBEAM-SMART-488, Toptica) for imaging the SYBR green stained phage. Images were acquired using customized SIM software described previously^20^.

### Optical infection assay

SYBR green stained T4 phage was added to overnight cultures of *E. coli* and *C. rodentium* in a 1:100 dilution ratio. The mix of phage and bacteria were incubated for 30 minutes at 37°C, before being passed through a Zeba spin desalting column (Thermo Scientific). The mixture was centrifuged at 3000 *g* for five minutes, and then resuspended in phosphate buffered saline (PBS, pH 7.4) for imaging. 2 μl of sample was deposited on a glass coverslip and an agarose pad was placed on top of the sample to prevent the bacteria from moving, as described previously. A glass coverslip was then placed on top of the agarose to prevent it from drying out. Images were acquired using SIM; the wavelength used for excitation was 488 nm (iBEAM-SMART-488, Toptica).

### T4 Phage adsorption assay

T4 phage adsorption assay was performed as described previously^21^. In this case, 10 ml of overnight *E. coli* culture was used as the host, infected with 1 μl of ~10^10^ PFU/ml T4 phage. As a no cell control, the same volume of phage was mixed with 10 ml of LB. The culture and the control were placed in an incubator at 37 °C, shaking at 150 rpm. Samples (100 μl) were taken at t = 0, 5, 15, 30 and 60 minutes and added to 900 μl LB and 20 μl NaHCO_3_ chloroform. The number of plaque forming units for each sample time point was determined by plaque assay as described in the phage propagation section above.

### OMV isolation from *E. coli* and *C. rodentium*

5 ml of LB broth was inoculated with *E. coli* cells and grown for 16-20 hours. 2 ml of the overnight culture was added to 200 ml of LB and allowed to incubate at 37 °C for ~3 hours until the OD600 of the culture was ~1.5. The cells were then centrifuged (4000 *g*, 4 °C, 15 minutes) to remove cell debris, and the supernatant was collected. The supernatant was further passed through a 0.22 μm filter. The outer membrane vesicles (OMVs) were then isolated by ultracentrifugation (140 000 *g*, 4 °C) for 3 hours (Beckman Coulter, Type 50.2 Ti Fixed-Angle Rotor) and the OMV pellets were resuspended in 250 μl of PBS supplemented with 2 mM MgCl_2_ solution. Finally, the OMV dispersion was centrifuged (16 000 *g*, 4°C) for 30 minutes to remove any final contaminants such as flagella. The supernatant was collected and re-suspended in 500 μl of PBS supplemented with 2 mM MgCl_2_ solution. The final OMV stocks were then stored at − 80 °C for further experiments.

For *C. rodentium* OMVs, the same protocol was followed except the culture was grown to an OD of ~ 2.0 prior to the same centrifugation and filtration steps.

### Preparation of POPC-PEG5kPE synthetic liposomes

POPC and PEG5kPE (Avanti) were mixed in a 99.5:0.5 molar ratio, with a nitrogen stream used to evaporate the chloroform, and the sample was further desiccated for one hour in a vacuum. The lipids were then hydrated in PBS supplemented with 2 mM MgCl_2_ to give a final lipid concentration of 4 mg ml^−1^. Single unilamellar vesicles were made by lipid extrusion through a 50 nm pore sized polycarbonate membrane, and samples were stored for up to two weeks at 4 °C.

### Dynamic Light Scattering (DLS)

DLS measurements were performed using a Zetasizer Nano S90 (Malvern Panalytical) configured with a 633 nm laser and a 90° scattering optic. The sample, one ml, was transferred into a disposable plastic cuvette, and three runs were taken for each measurement. The intensity of the scattered light is used by the Zetasizer software to determine three main parameters: Z average (d.nm) which measures the average size of the particle distribution; count rate (kcps) which counts the number of photons detected per second and is related to the concentration and quality of the sample; polydispersity index (PDI) which provides a measure for the heterogeneity of the particle size distribution.

### Nanoparticle Tracking Analysis (NTA)

Nanoparticle Tracking Analysis (NTA) was carried out using a Nanosight NS500 (Malvern Panalytical) fitted with an Electron Multiplying Charged Couple Device (EMCCD) camera configured with a 522 nm laser. Prior to analysis, samples were diluted (1:500) in PBS. 5 × 60 second videos were recorded for each sample analysed, with a temperature range of 20.8 – 21.5°C and a camera level of 15. NTA 3.2 software was used to analyse the data with a detection threshold of 5.

### Transmission Electron Microscopy (TEM)

For imaging OMVs, 10 μl of sample (concentration 10^10^ particles/ml) was negatively stained with 1% (w/v) uranyl acetate solution for two minutes at room temperature before being visualised with a Tecnai G2 80-200 keV transmission electron microscope, operating at 200 keV with images recorded with a bottom-mounted AMT CCD camera. For T4 phage imaging, 10 μl of sample (concentration 10^9^ PFU ml^−1^) was used whilst for T4 and OMV combined imaging, phage (~ 10^9^ PFU ml^−1^) was mixed with *E. coli* OMVs (~10^10^ PFU ml^−1^) in a 1:1 ratio, and left to incubate at room temperature for 20 minutes before the negative staining procedure was conducted.

### Formation of supported lipid bilayers on glass coverslips

Glass coverslips (Academy, 22×40 mm, 0.16-0.19 mm thick) were first cleaned with acetone and isopropanol in a 1:1 ratio. 100 μl of ~10^10^ OMV particles ml^−1^ was added to the glass slide and allowed to incubate for 20 minutes before washing twice with PBS solution to remove excess unadhered OMVs. 100 μl of POPC-PEG5kPE liposomes were then added for one hour to induce rupturing of the OMVs. The well was then washed again twice with PBS; the SLBs were then kept in PBS solution for imaging.

### Characterization of SLBs using fluorescence recovery after photobleaching (FRAP)

Prior to analysis by fluorescence recovery after photobleaching (FRAP), OMVs must be fluorescently labelled. This was achieved by adding 1 μl of R18 dye (Invitrogen) to 200 μl of OMV stock, and sonicating for 15 minutes. A G25 spin column (GE healthcare) was used to remove unbound/excess R18 by centrifugation at 1500 *g* for 3 minutes at room temperature. Lipid bilayers were formed using the protocol outlined above.FRAP measurements were conducted using an inverted Zeiss LSM800 confocal microscope with a 10x objective lens. A 30 μm diameter bleaching spot was made, and recovery of the fluorescence intensity of this spot was measured over time relative to a 50 μm diameter reference spot. The data were analyzed using MATLAB, and the fluorescence recovery was modelled using a modified Bessel function as previously described^22^. The model fit was used to extract the diffusion coefficient (D) according to the equation D = r^2^/4*τ* where *r* is the radius of the photobleached spot and *τ* is the characteristic diffusion time. The fit was also used to extract the mobile fraction (MF) according to the equation (I_E_ - I_0_) / (I_I_ - I_0_), where I_E_ is the final postbleach intensity value, I_0_ is the first postbleach intensity value and I_I_ is the initial prebleach intensity value.

### SIM imaging of phage SLB interactions

SLBs were formed in the manner described above, where the lipids were stained with R18. SYBR green stained phage (concentration ~ 10^10^ PFU ml^−1^) were added to the SLBs (*E. coli/ C. rodentium/* POPC-PEG5kPE) and incubated for 20 minutes, washed with PBS before imaging using SIM. To acquire SIM images, a ×60/1.2 NA water immersion lens (UPLSAPO 60XW, Olympus) focused the structured illumination pattern onto the sample, and the same lens was also used to capture the fluorescence emission light before imaging onto an sCMOS camera (C11440, Hamamatsu). The wavelengths used for excitation were 561 nm (OBIS 561, Coherent) for the lipid bilayers and 488 nm (iBEAM-SMART-488, Toptica) for the SYBR green stained phage.

### QCM-D measurements

QCM measurements were performed using a Q-sense analyzer (QE401, Biolin Scientific). Piezoelectric silicon sensors were used for all the experiments. First the frequency and dissipation of energy signals were stabilized in PBS and the different lipid vesicles stocks in PBS were pumped into the chamber with a constant flow rate of 70 μL min^−1^ controlled by a peristaltic pump. All vesicles were given enough time to adsorb on the surface of the crystal. After that PBS 1X solution was used to wash unbound vesicles. When a stable frequency signal was established, phage was pumped in the chamber at a constant flow rate of 70 μl min^−1^ and incubated for about an hour on both synthetic lipids-functionalized sensors and outer-membrane lipids functionalized QCM sensors. Finally, the sensors were rinsed with PBS. The mass reported in figure 3 was extracted from raw frequency and dissipation of energy data (please add supporting figure if there is any) using both the Young-Voigt viscoelastic model and Sauerbrey equation^23^. Q-tools, D-find and Q-soft software were used for the modelling the raw data.

### Preparation of PEDOT:PSS mixture

PEDOT:PSS dispersion (Heraeus) was mixed with 5% (v/v) ethylene glycol (EG) and 0.5 % (v/v) dodecylbenzenesulfonic acid (DBSA) in order to enhance film formation and conductivity. 1% (v/v) 3-glycidoxypropyltrimethoxysilane (GOPS), which is a polymer crosslinking agent, was then added and the final mixture was sonicated for 10 minutes and sequentially filtered through a 0.8 μm and a 0.45 μm syringe filter prior to use.

### Microelectrode array device fabrication

PEDOT:PSS microelectrode arrays were fabricated using two different device architectures with both parylene C and silicon oxide used as electrode insulation layers. Parylene C devices are designed into arrays with circular electrodes with 450 μm in diameter or 200 μm by 200 μm square electrodes. To fabricate the devices, 4 inch glass wafers first were cleaned by sonication in acetone and then isopropanol for 15 minutes. The wafers are rinsed with DI water and baked 15 minutes at 150 °C. To pattern for contact tracks, a negative photoresist, AZ nLOF2035 (Microchemicals GmbH) was spun on the glass wafer with 3000 rpm for 45 s and exposed with UV light using mask aligner (Karl Suss MA/BA6). The photoresist was developed in AZ 726 MIF developer (MicroChemicals) developer for 28 s. Ti (5 nm)/Au (100 nm) layer as conductive tracks was deposited by e-beam evaporation on top of the wafer and the Ti-Au metal layer was lifted-off by soaking in Ni555 (Microchemicals GmbH) overnight. Prior to the deposition of 2 μm layer (sacrificial layer) of parylene C ((SCS), the wafer was soaked with 3% A174 (3-(trimethoxysilyl)propyl methacrylate) in ethanol solution (0.1% acetic acid in ethanol) for 60 s to promoteparylene C adhesion onto the wafer. An anti-adhesive layer of Micro-90 in DI water (2% v/v solution) was spun (1000 rpm for 45 s), and then the second layer of 2 μm parylene C (SCS) was deposited. A layer of positive photoresist AZ 10XT (Microchemicals GmbH) was spun at 3000 rpm for 45 s, exposed to UV, and developed in AZ 726 MIF developer (MicroChemicals) for 6 minutes to pattern electrode areas. Reactive ion etching (Oxford 80 Plasmalab plus) opened the window for deposition of Clevios PH500 PEDOT:PSS (Heraeus). The PEDOT:PSS mixture (prepared as described above) was spin coated at 3000 rpm for 45 s. The device was baked at 90 °C for 1 minute, and the sacrificial parylene C layer was peeled off. Finally, the sample was baked at 130 °C for 1 hour.For devices fabricated using the silicon oxide method, glass wafers were first cleaned by heating in 9:1 sulfuric acid/hydrogen peroxide for 20 min at 120 °C. Metal contacts (50 nm Au between two 5 nm Ti adhesion layers) were e-beam evaporated and photolithographically patterned using a lift-off process. 230 nm SiO2 insulation layer was deposited with chemical vapor deposition and Au contacts were photolithographically patterned and then exposed using inductively coupled CHF3 plasma reactive ion etching. Wafers were treated with O2 plasma and then PEDOT:PSS layer was deposited via spin-coating the PH 1000 (Ossila) dispersion containing 5% v/v ethylene glycol and 1% v/v (3-glycidyloxypropyl)trimethyoxysilane at 2000 RPM for 2 min and baked at 140°C for 30 min. 100 nm Ge hard mask was deposited using e-beam evaporation. Ge layer and underlying PEDOT:PSS were patterned using photolithography and inductively coupled plasma reactive ion etching with CF4 and O2 respectively. Devices were soaked in deionized water for 48 hr to oxidize and remove the Ge layer. Electrodes were 500 × 500 μm square electrodes.

### EIS measurements

EIS was performed using a potentiostat (Autolab PG-STAT204) in a three-electrode configuration with Ag/AgCl and Pt electrodes being used as the reference and counter electrodes respectively. Each PEDOT:PSS coated gold electrode in a single array was sequentially used as the working electrode. The AC current was recorded within the frequency range 50 – 100 000 Hz, with 10 data points per decade (equally spaced on a logarithmic scale). An AC voltage of 0.01 V and a DC voltage of 0 mV versus OCP were applied. For all experiments, LB was used as the electrolyte. Measurements were recorded for the baseline (ie. no bilayer), after bilayer formation (as described above) and after phage (concentration ~ 10^10^ PFU ml^−1^) was incubated with the bilayer for 20 minutes. Data was collected and analysed using NOVA 2.1.3 software (Metrohm Autolab).

## RESULTS AND DISCUSSION

We first characterized the T4 phage particles using both AFM and TEM imaging (Fig 1bi, bii). The total length of the phage was 239 ± 13 nm, with the hexagonal head having length and width of 119 ± 9 nm and 105 ± 17 nm respectively. These dimensions were in agreement with previously reported values for T4 phage^24^. *E. coli* and *C. rodentium* are both members of the attaching and effacing (A/E) family of bacteria and are genetically very similar, sharing large parts of their genomic sequences^25,26^. Even though these two types of bacteria are similar, T4 phage was only able to infect *E. coli* and not *C. rodentium* as shown by SIM imaging of infectivity assays, since T4 phage particles could be stained with SYBR green (SFig 1). After incubation with stained T4 phage (Fig 1ci, 1cii), a significant difference was seen in the fluorescence of the two types of bacteria (558±80 and 360±5 AU for *E. coli* and *C. rodentium* respectively, P = 0.0006). Further evidence that T4 phage was only able to infect *E. coli* and not *C. rodentium* was shown by plaque assays, thus providing a means to exploit the difference in specificity to validate the eventual screening platform (Fig 1d). The area of clearance, or plaques, in the *E. coli* plate evidenced the phage infection. We investigated the kinetics of the T4 interaction with *E. coli* cells by conducting a phage adsorption assay (Fig 1e, SFig 2). By counting the number of plaques, and therefore measuring the PFU ml^−1^ over time (indicative of free, unbound phage), we observed the concentration significantly decreased from 2.4 ± 0.4 (x 10^4^ PFU ml^−1^) to 0.9 ± 0.1 (x 10^4^ PFU ml^−1^) after 15 minutes (P = 0.018). This was in contrast to the control case, where the concentration did not significantly change between 0 and 15 minutes (concentrations were 2.8 ± 0.1 and 2.7 ± 0.1 (x 10^4^ PFU ml^−1^) respectively. Combining all this information, we concluded that that the choice of bacteria and phage were appropriate for moving forward with developing the SLB screening platform.

The process of generating *E. coli* OM SLBs is depicted in Figure 2a. Briefly, OMVs isolated from an *E. coli* culture were induced to rupture and fuse, producing a complete SLB with the addition of POPC-PEG fusogenic liposomes. The presence of OMVs was confirmed using nanoparticle tracking analysis (NTA), a technique that measures the size distribution of particles based on their diffusivity^27^. The average size of the particles was 186.6 ± 18.3 nm, in line with the expected size of *E. coli* OMVs^28^. TEM was used to image the OMVs in the sample (SFig 3a), verifying their morphology in line with previous images of OMVs^29^. The quality of the resulting SLB after vesicle fusion was evaluated using fluorescence recovery after photobleaching (FRAP). This technique relies on fluorescently staining OMVs with rhodamine-18 (R18) in the SLBs and then observing how, when a spot in the bilayer is bleached, fluorescence intensity recovers over time due to the lipid lateral mobility. As shown by figure 2b, the *E. coli* SLB fluorescence recovered over time, indicating that the bilayer was complete and mobile. The diffusion coefficient (D) and mobile fraction (MF) values were 0.59 ± 0.04 μm^2^/s and 0.84 ± 0.04 respectively, similar to the values obtained by Hsia *et al*. for OM SLBs on glass^13^. Since *C. rodentium* also produce OMVs^30^, a crucial aspect of the screening procedure was the ability to form SLBs using the same process as for *E. coli* SLBs (Fig 2a). We confirmed the isolation of *C. rodentium* OMVs using NTA (Fig 2ci) and TEM (SFig 3b) with the OMVs having an average size of 134.6 ± 4.9 nm. SLBs were also generated, with the FRAP data showing the bilayers to have D and MF values of 0.39 ± 0.03 μm^2^/s and 1.02 ± 0.02 respectively.

**Figure 2.**
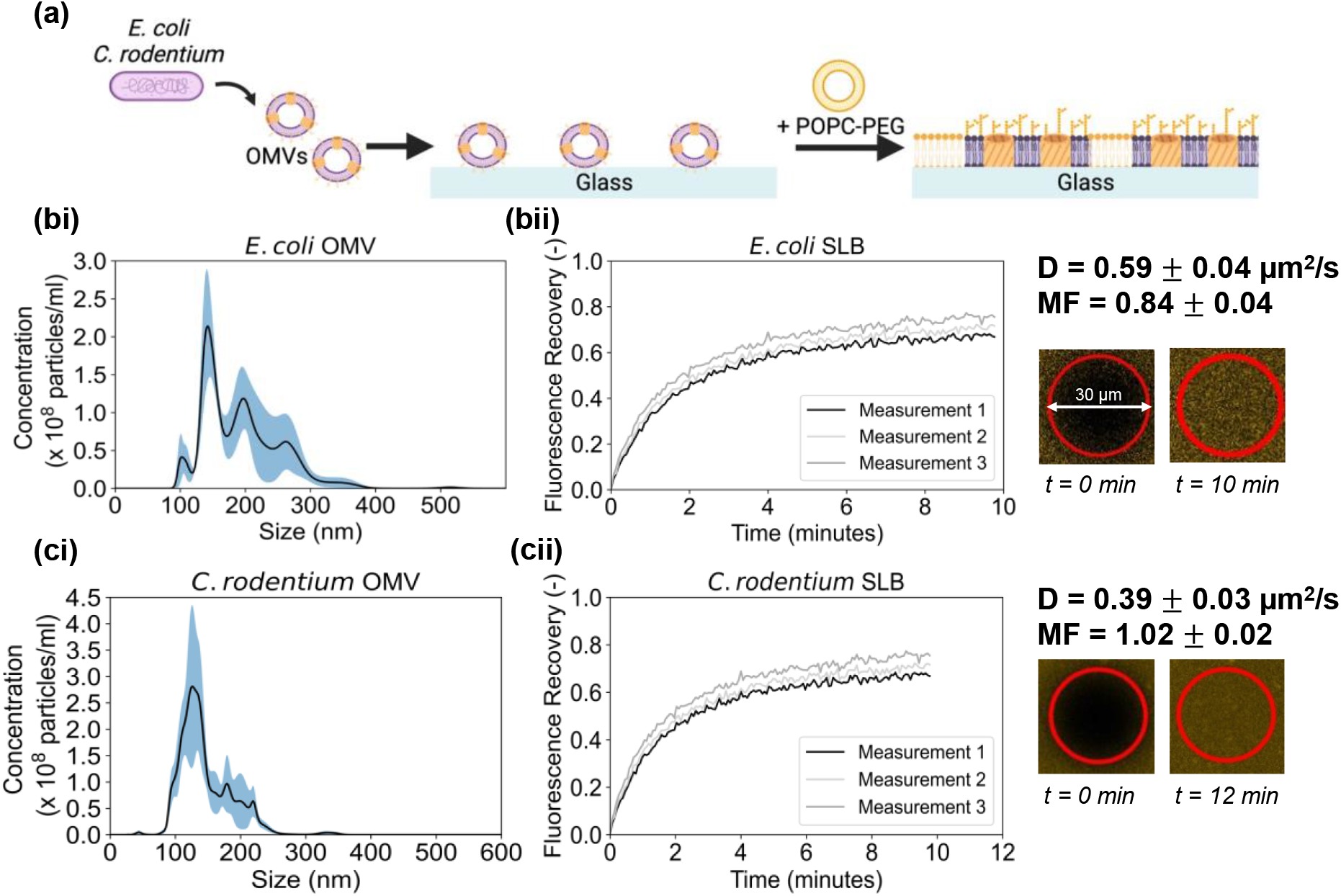
Formation of SLBs derived from bacterial OMVs. (a) Schematic showing the process of forming OM SLBs using *E. coli* OMVs. (bi) NTA characterization of the *E. coli* OMVs, with the main peak at 143 nm and the average size being 186.6 ± 18.3 nm. The blue shading represents the standard error of the mean. (bii) FRAP data for the *E. coli* OM SLB, showing the recovery of the bleached spot over time due to the mobility of the lipids. The corresponding diffusion coefficient and mobile fraction values are 0.59 ± 0.04 μm^2^/s and 0.84 ± 0.04 respectively. Diameter of the bleached spot is 30 μm. (ci) NTA characterization of the *C. rodentium* OMVs, showing a mean peak at 142 nm and the average size at 134.6 ± 4.9 nm. The blue shading represents the standard error of the mean. (cii) FRAP data for the *C. rodentium* SLB, where the diffusion coefficient and mobile fraction values are 0.39±0.03 μm^2^/s and 1.02±0.02 respectively.

Having established that SLBs could be generated using the two strains of bacterial OMVs, T4 phage binding to the *E. coli* and *C. rodentium* SLB systems was evaluated. SYBR green stained T4 phage incubated with R18 stained SLBs were imaged by SIM. With *E. coli* SLBs, spots of green fluorescence can be observed, indicating that the phage particles could interact with the SLB due to the presence of LPS and OmpC – the two components required for initial T4 phage binding at the start of the infection process in whole cells^31^ (Fig 3ai). Conversely, no binding was observed with the *C. rodentium* SLB (SFig 4); this shows the high specificity of the binding interaction since even though the bacterial strains are closely related, the outer membrane did not allow for phage binding. The fluorescence quantification verified this – the fluorescence in the 488 nm wavelength region was ~3 times higher for the *E. coli* SLBs compared to the *C. rodentium* SLBs (Fig 3aii). POPC only SLBs were used as an additional negative control showing a similar lack of T4 phage binding (SFig 5).

**Figure 3.**
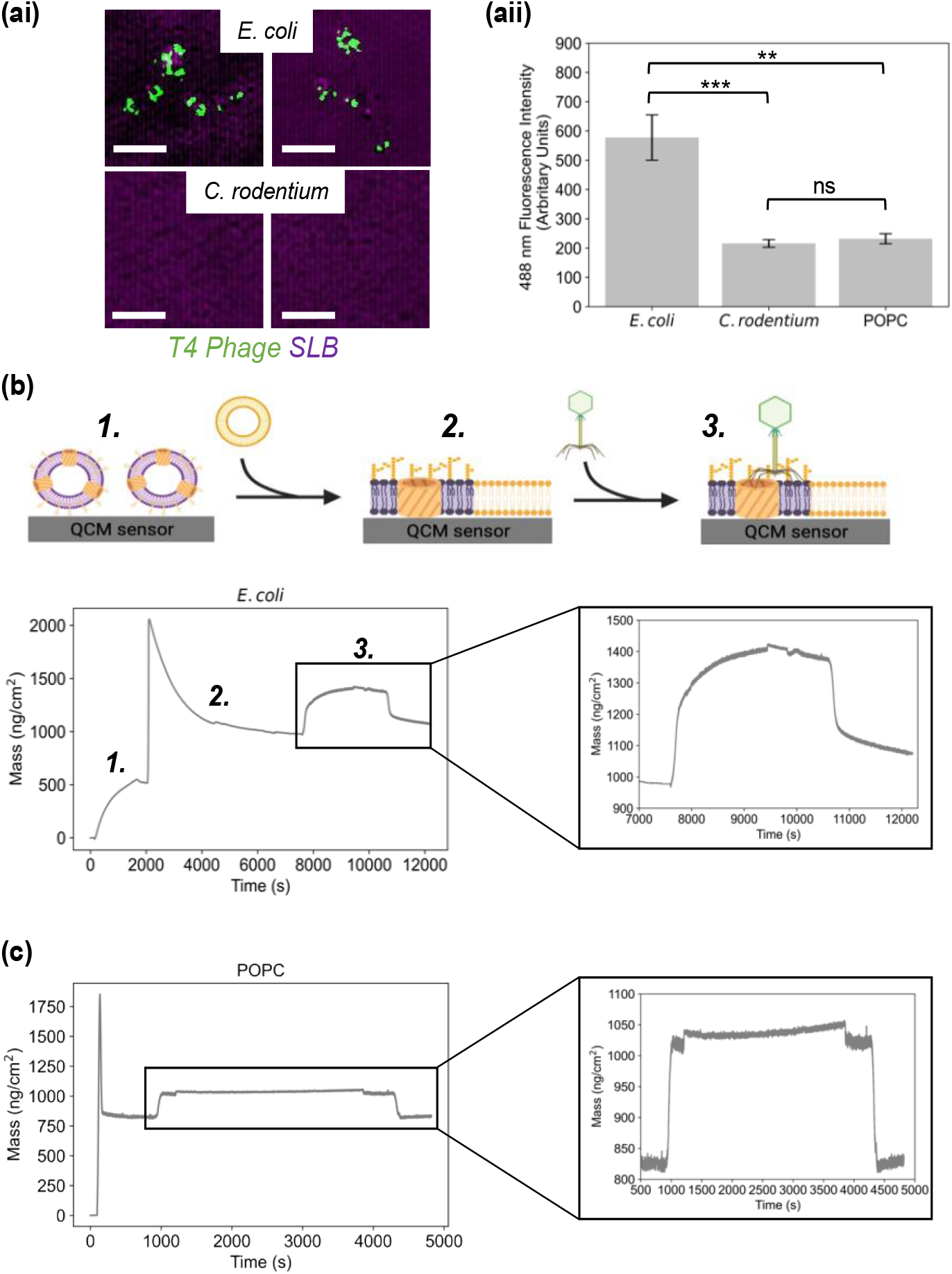
Optical and QCM-D screening of T4 phage interactions with SLBs. (ai) Panel of sim images taken from *E. coli* and *C. rodentium* OM SLBs (stained with R18) incubated with SYBR green stained T4 phage. Scale bar = 2 μm. (aii) Measurement of the fluorescence intensity in the 488 nm wavelength region (measured in arbitrary units) for the 3 types of bilayers, including POPC SLB, after phage incubation. Error bars represent the standard error (two-way analysis of variance; *P≤0.05; **P≤0.01; ***P≤0.001; n = 6). (b) QCM-D measurements for phage interaction with SLBs. Schematic of the sequence of events in the formation of the OM SLB and subsequent phage addition is shown. (1) Addition of OMVs, (2) Addition of POPC-PEG and SLB formation, (3) Addition of phage. Graph below shows mass changes over time due to phage addition to the OM SLB. Phage is added at ~ 7800 s and washing commences at ~ 10600 s. (c) Corresponding QCM-D measurements for the control POPC SLB and subsequent phage addition. Zoom in of the portion of the graph related to the phage addition. Phage is added to the system at ~ 1000 s and washing commences at ~ 4400 s.

QCM-D is a technique that has been used extensively to monitor SLB formation and subsequent interactions and so we used it to verify the phage SLB interaction^32^. This technique relies on measuring changes in frequency and dissipation at various overtones (these correspond to various penetration depths of the signal) on a SiO_2_ sensor. The changes in frequency (Δf) relate to changes of mass absorbed on the sensor; a decrease in Δf denotes an increase in mass. Alternatively, changes in dissipation (ΔD) are related to a change in the rigidity where an increase in ΔD denotes a decrease in rigidity. We first monitored the formation of the *E. coli* OM SLB using QCM-D. Focusing first on the Δf over time (SFig 6ai), we observed that after OMVs were added, there was a decrease of ~20 Hz due to the mass increase of the absorbed OMVs. This Δf is in line with previously reported values for Gram-negative bacteria OMVs adsorbed to the SiO_2_ sensor^14^. After addition of the POPC-PEG liposomes, there appeared a sharp drop in Δf and concurrent increase in ΔD (SFig 6aii) as the liposomes adsorbed to the surface. These were followed by an increase in Δf and drop in ΔD, a signature that is associated with a release of coupled-mass water seen in these systems as the vesicles rupture and fuse to form the SLB^13,14,33^. Figure 3b shows how the mass of the adsorbed material on the sensor was extracted from the Δf and ΔD data presented in the supplementary information. At ~ 8000 s, the T4 phage was added and an increase in mass was observed, suggesting that the phage bound to the surface of the SLB. At ~ 10600 s, when buffer was washed over the sensor, there was a mass difference of ~ 100 ng cm^−2^ after washing. This change in mass was indicative of permanent and specific phage binding – since the mass of a single phage virion is 194 MDa (when the capsid head is filled with DNA), this mass increase corresponded to the binding of ~ 3.11 × 10^8^ phage cm^−2^. To further confirm the specificity of T4 phage binding, a control QCM-D measurement was conducted with a POPC bilayer. The rapid decrease in Δf followed by an increase (occurring simultaneously with the opposite trends in ΔD) in the signal indicated that the POPC-PEG liposomes were adsorbed to the sensor surface, and then rapidly ruptured and fused to form an SLB with the associated release of coupled-mass water (SFig 5bi, 5bii). The Δf signal stabilized at - 28 Hz which is in line with synthetic bilayer formation monitored by QCM-D on SiO_2_ sensors^34^. Figure 3c illustrates the mass changes extracted from the Δf and ΔD data. At ~1000 s, phage was added to the system and an increase in mass was observed from ~ 820 ng/cm^2^ to ~ 1030 ng/cm^2^. However, when the sensor was washed with buffer in the same manner as for the *E. coli* SLB system, the mass dropped back to the pre phage incubation level of ~ 820 ng/cm^2^. This indicated that the phage could not bind to the SLB specifically, in contrast to what was seen in the *E. coli* SLB case. The QCM-D experiments thus complemented the findings of the SIM experiments and expanded upon them to an extent since not only the phage-SLB interactions but also the bilayer formation itself were monitored to a high degree of precision.

A fundamental part of this study was the creation of a platform that provided a quick and quantitative evaluation of phage interaction with a given bilayer system. EIS can be used to measure the properties of an SLB integrated on a MEA device. EIS measures the compound effect of resistance and capacitance properties of an SLB by applying an alternating voltage over a frequency range and measuring the corresponding impedance (Z), which is a measure of resistance to current flow in the system. By decoupling the real part and imaginary part of SLB impedance, the membrane resistance and capacitance can be extracted. In our setup, when integrating the SLB with PEDOT:PSS coated MEA devices, we used a well-established equivalent circuit in which the bilayer is modelled as a resistor and capacitor in parallel^35,36^ (Fig 4a). The complex Z consists of an imaginary and real part; when these are plotted against each other for each frequency, a Nyquist plot is generated. The Nyquist plot is a useful way of expressing EIS data since the width of the semi-circle portion of the graph denotes the membrane resistance.

**Figure 4.**
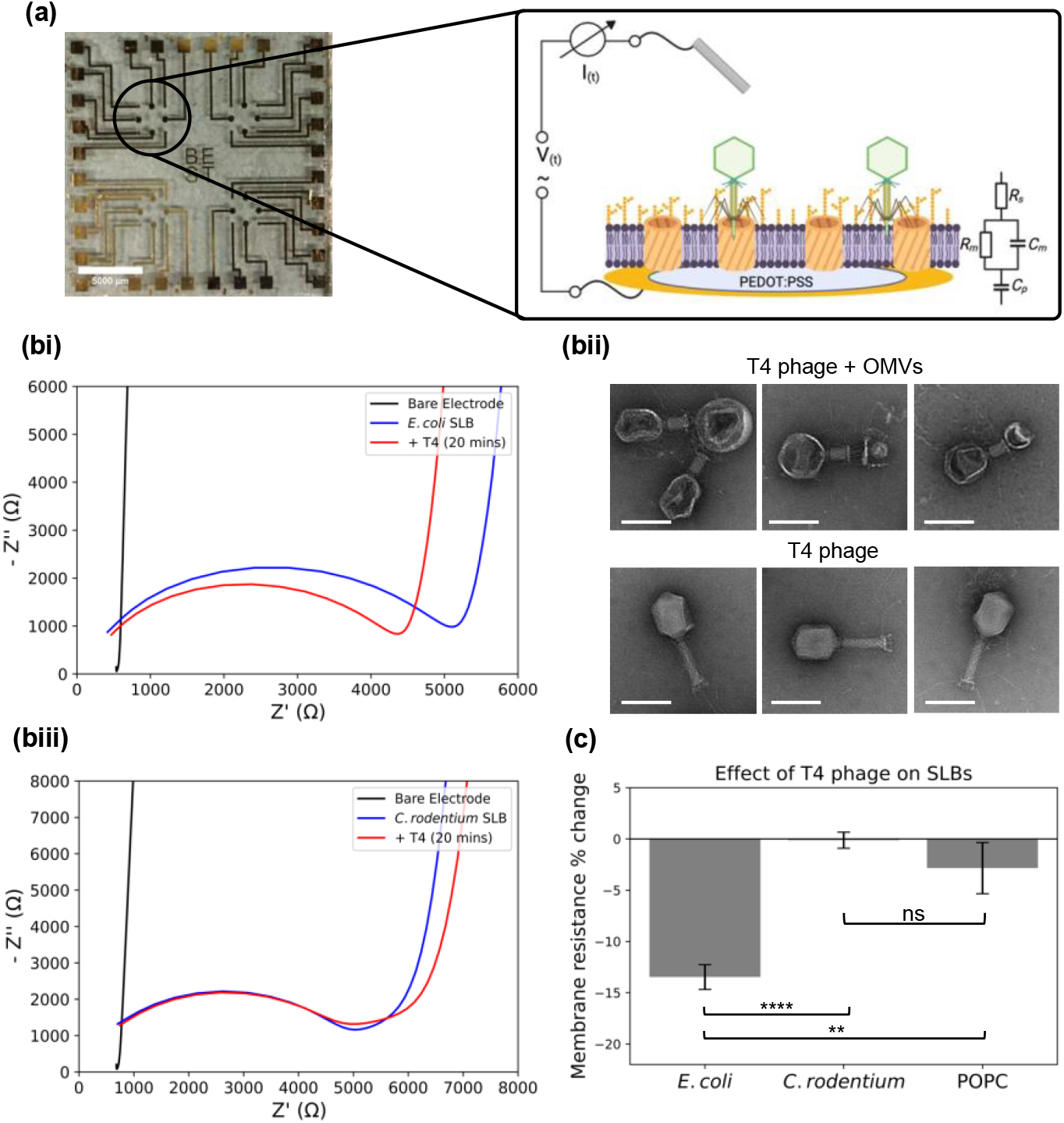
Electrical screening of phage interaction with SLBs. (a) Image of a MEA used for the EIS measurements. Scale bar = 5 mm. Each array consists of eight PEDOT:PSS coated electrodes upon which the SLB is formed. The impedance of the SLB is modelled using the equivalent circuit in the inset shown. (bi) Nyquist plot from a representative EIS recording for T4 phage incubated with *E. coli* SLBs. Measurements were taken before the SLB was formed, after the SLB formation and after phage incubation. The decrease in the impedance is hypothesized to be the result of the phage particles binding and initiating pore formation in the OMV component of the SLB due to phage tail contraction and tube penetration. (bii) TEM images of T4 phage, comparing the morphology with and without interaction with *E. coli* OMVs. Scale bar = 100 nm. (biii) Nyquist plot from a representative EIS recording for T4 phage incubated with *C. rodentium* SLBs. Measurements were taken before the SLB was formed, after the SLB formation and after phage incubation. (c) Membrane resistance % change (ie. change in extracted membrane resistance before and after phage incubation) on each electrode for three types of SLBs (*E. coli* SLB, *C. rodentium* SLB, POPC SLB). Error bars represent the standard error (two-way analysis of variance; *P≤0.05; **P≤0.01; ***P≤0.001; n = 3).

When EIS was used to monitor the *E. coli* SLB formation on the MEA, the characteristic semi-circle shape on the Nyquist plot appeared, denoting the presence of an additional RC element, indicating the presence of an SLB (Fig 3bi). The Bode plot also showed the presence of this insulating barrier (SFig 6a). The extracted resistance for the SLB was 251.6 ± 34.1 Ω.cm^2^, in agreement with previously reported values for SLBs containing mammalian cell components^37^. When T4 phage was added to the bilayer, a 14 % reduction in the membrane resistance was observed (217.5 ± 30.1 Ω.cm^2^) – we hypothesized that this was due to the interaction process occurring between the phage particles and the outer membrane components on the SLB surface. When T4 interacts with a cell, the initial process involves the adhesion of the phage to the outer membrane surface, followed by sheath contraction and subsequent tail tube penetration through the outer membrane^38^. This penetration by the tail tube and effective pore formation would explain the reduction in resistance observed here – as the phage particles interact with the *E. coli* SLB, the attachment and penetration processes are initiated, leading to increased ion flow through the SLB and subsequent drop in membrane resistance. To investigate whether intact OMV components could trigger this membrane penetration process, we used TEM to image the phage bound to *E. coli* OMVs since these are the components of the SLB that would interact with the phage (Fig 4bii). The difference between phage particles bound and unbound to OMVs was stark – the bound phage exhibited contracted tails (about half of the uncontracted length), with this contraction driving the penetration of the tail tube through the OMV membrane^39^. Moreover, the morphology of the capsid head was altered, which we postulate to be associated with the release of DNA through the tail tube. This supported the notion that the drop in impedance seen in the EIS readings was due to the phage tail tube penetration of the bilayer taking place.

*C. rodentium* SLBs were also generated on MEAs and probed by EIS. The Nyquist (Fig 4biii) and Bode (SFig 6b) plots confirmed the formation of the SLB, with the extracted membrane resistance being 230.2 ± 46.7 Ω.cm^2^ which was very similar to the resistance observed for the *E. coli* SLB. Crucially, when phage was added here, a 1% decrease in membrane resistance was observed (227.9 ± 44.9 Ω.cm^2^), indicating that T4 phage was unable to attach and subsequently penetrate these bilayers due to the absence of the necessary outer membrane components. Finally, POPC SLBs were used as an additional control (SFig 6ci, 6cii). EIS measurements showed the bilayer resistance to be 481.4 ± 168.8 Ω.cm^2^ initially and 447.0 ± 149.7 Ω.cm^2^ after phage addition. Due to electrode-to-electrode variation in bilayer quality, it was important to measure relative changes on each electrode individually to meaningfully compare between data sets. When relative changes in membrane resistance were measured, it became clear that the decrease in membrane resistance upon phage incubation was found only in the *E. coli* SLB case (Fig 4biv). Here, the membrane resistance decreased by 13.5 ± 1.2 % whilst it decreased by only 0.1 ± 0.8 % and 2.8 ± 2.5% in the *C. rodentium* and POPC SLBs respectively. Therefore, the difference in membrane resistance changes between the three SLB cases provides a quantitative screening mechanism for detecting phage – SLB interactions.

The results here describe a multiparametric, quantitative readout of T4 phage interacting with OM SLBs. T4 phage belongs to the *myoviridae* family of phage characterised by their long contractile tails. *Podoviridae*, exemplified by T7 phage, have short non contractile tails and therefore exhibit a different mechanism of interaction with outer membranes and subsequent pore formation^39,40^. On the other hand, the *siphoviridae* family of phage (eg. λ phage) have long non contractile tails and contain a central tape measure protein used to dictate tail length as well as to facilitate DNA injection through the outer membrane^41^. We therefore envisage this platform to measure the interaction not only with *myoviridae* and OM SLBs, but also with these other phage families to analyse whether the results we see depend on the mechanistic variation of phage infection. Perhaps there is a certain electrical signature that distinguishes these various mechanisms, in a similar manner to how EIS can be used to distinguish between different mechanisms of antibiotic interaction with membranes^18^.

In terms of advancing phage therapy, there is considerable interest on how variations on both the tail structures of phage, as well as the receptors present in the outer membrane, help or hinder an infection event. Recently, Zeng and Salmond were able to increase the host range specificity of lambda phage by expressing its receptor LamB in three different bacterial genera ^42^. The OM SLB platform could act as a complementary technique to those used in this study, due to the ease of generating OMVs derived from these various mutant bacteria. In a similar manner, rational engineering of the proteins in the phage tails responsible for receptor binding, the so-called phage receptor binding proteins (RBPs), has been used to programme the host specificity of the phage. For instance, Yehl *et al*. used site directed mutagenesis to create a T3 phage library with different RBP sequences and thus different host range specificities^43^. The SLB platform could help with evaluating whether a given phage tail mutation increases or decreases the propensity of the phage to interact with bacterial membranes via interpretation of the optical, surface sensitive and electrochemical interaction signatures described in this paper.

Bacteria produce structures as part of their innate immune defence systems that are morphologically very similar to phage tail structures. These so called tailosins are produced by bacteria to destroy closely related strains, and their mechanism of action relies on compromising the bacterial cell membrane^44^. For instance, S5 pyocins destroy *Pseudomonas aeruginosa* strains by delivering a pore forming complex across the OM^45^. By measuring interactions between tailocins and OM SLBs, this would provide information on the potential of the structures as antimicrobials against a given pathogenic bacteria. Similarly, defensins are phage tail like structures produced by mammalian immune system cells to destroy pathogenic bacteria. Studies on alpha defensins have shown their pore forming mechanism of action in synthetic lipid bilayers^46^, and we hope to demonstrate the ability of our OMV derived SLB system to monitor the disruption effects of novel defensins, and therefore extend the field of defensin research as an alternative to current antimicrobial strategies.

## CONCLUSION

In conclusion, we have shown that SLBs containing the naturally occurring components of bacterial outer membranes provide a platform for specifically detecting bacteriophage interactions. After first confirming two types of bacteria, namely *E. coli* and *C. rodentium*, that can be used as the basis for a T4 phage screening platform, we showed that OM SLBs using OMVs isolated from these two bacteria can be generated. Optical imaging with SIM revealed that T4 was only able to interact with the *E. coli* SLB, and this interaction was further quantified using the highly sensitive QCM-D monitoring. The culmination of the study was the electrochemical monitoring of the differential phage interaction with the three types of SLBs, thus providing a quick and quantitative readout of the interaction (or lack thereof). In addition to this, the MEA platform has the potential to be integrated with a multiplexed, microfluidic setup which could prove pivotal in developing high throughput phage screening methods. Although in this study we sequentially use optical, surface sensitive and electrical techniques, future work will be based on simultaneous monitoring on a single multipurpose device. Overall, the study outlined here is the first time SLBs have been used to interact with phage, and the setup could prove important in easily and quantitively determining hereto unknown phage-bacteria interactions. This will help accelerate the advent of phage therapy - a crucial avenue that needs to be explored further if our fight against antibiotic resistance is to prove successful.

## Supporting information

supplementary_phage_Karan_Bali

## ASSOCIATED CONTENT

### Supporting Information

Supporting information available: The following files are available free of charge: supplementary_acs_sensors_phage_KB.pdf. It contains the plates used for the adsorption assay and TEM images of OMVs used to make SLBs. In addition, it contains a panel of SIM images for the fluorescent screening of phage interactions, as well as extra QCM-D and EIS data.

## AUTHOR INFORMATION

### Author Contributions

KB grew and isolated OMVs, propagated phage samples and performed FRAP, EIS and plaque and adsorption assay measurements. GS and RM (Monson) provided bacterial glycerol stocks and T4 phage. ZL, JT and AS (Salleo) designed and fabricated PEDOT:PSS microelectrode arrays. AS (Savva) performed QCM-D measurements and analysed these data. KB, CFK and IM performed SIM measurements. RM (McCoy) performed NTA measurements. KB analysed the data, wrote and revised the manuscript with RM (Monson) and RMO. RM (Monson) and RMO supervised the study and provided funding support. All authors approved the manuscript prior to submission.

## ACKNOWLEDGMENT

All figures were created using BioRender.com. K.B. was funded by an Engineering and Physical Sciences Research Council (EPSRC)-DTP PhD studentship (project 2266415). R.M. acknowledges funding from the EPSRC Cambridge NanoDTC under award EP/S022953/1. Part of this work was performed at the Stanford Nano Shared Facilities (SNSF), supported by the National Science Foundation under award ECCS-2026822. Part of this work was performed in part in the nano@Stanford labs, which are supported by the National Science Foundation as part of the National Nanotechnology Coordinated Infrastructure under award ECCS-2026822.We acknowledge the Cambridge Royce facilities grant EP/P024947/1 and Sir Henry Royce Institute - recurrent grant EP/R00661X/1 for EQCM-D facility at Maxwell Centre, University of Cambridge. A.S. acknowledges funding from the European Union’s Horizon 2020 research and innovation program under the Marie Skłodowska-Curie grant, MultiStem (No. 895801). C.F.K. acknowledges funding from the EPSRC (EP/H018301/1 and EP/L015889/1), the Wellcome Trust (089703/Z/09/Z and 3-3249/Z/16/Z), the Medical Research Council (MR/K015850/1 and MR/K02292X/1), MedImmune, and Infinitus (China). I.M. acknowledges funding from the Royal Society (URF/R1/221795). R.M.O. acknowledges funding for this project sponsored by the Defense Advanced Research Projects Agency (DARPA) Army Research Office and accomplished under Cooperative Agreement W911NF-18-2-0152. The views and conclusions contained in this document are those of the authors and should not be interpreted as representing the official policies, either expressed or implied, of DARPA or the Army Research Office or the U.S. Government. The U.S. Government is authorized to reproduce and distribute reprints for Government purposes notwithstanding any copyright notation herein.

## For Table of Contents Only

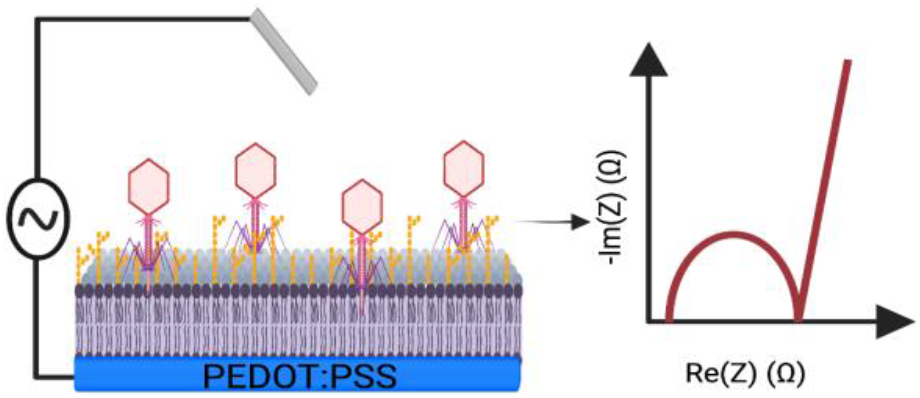

