## supplementary_phage_Karan_Bali for "Multiparametric sensing of outer membrane vesicle-derived supported lipid bilayers demonstrates the specificity of bacteriophage interactions"

SUPPLEMENTARY MATERIAL

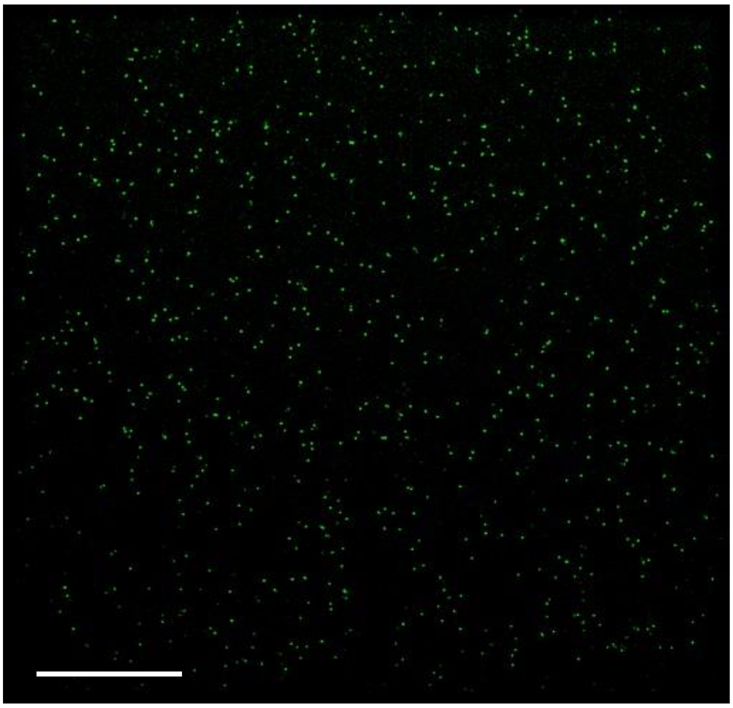

SFig 1. SIM imaging of SYBR green stained T4 phage on a glass slide. Scale bar = 10 μm.

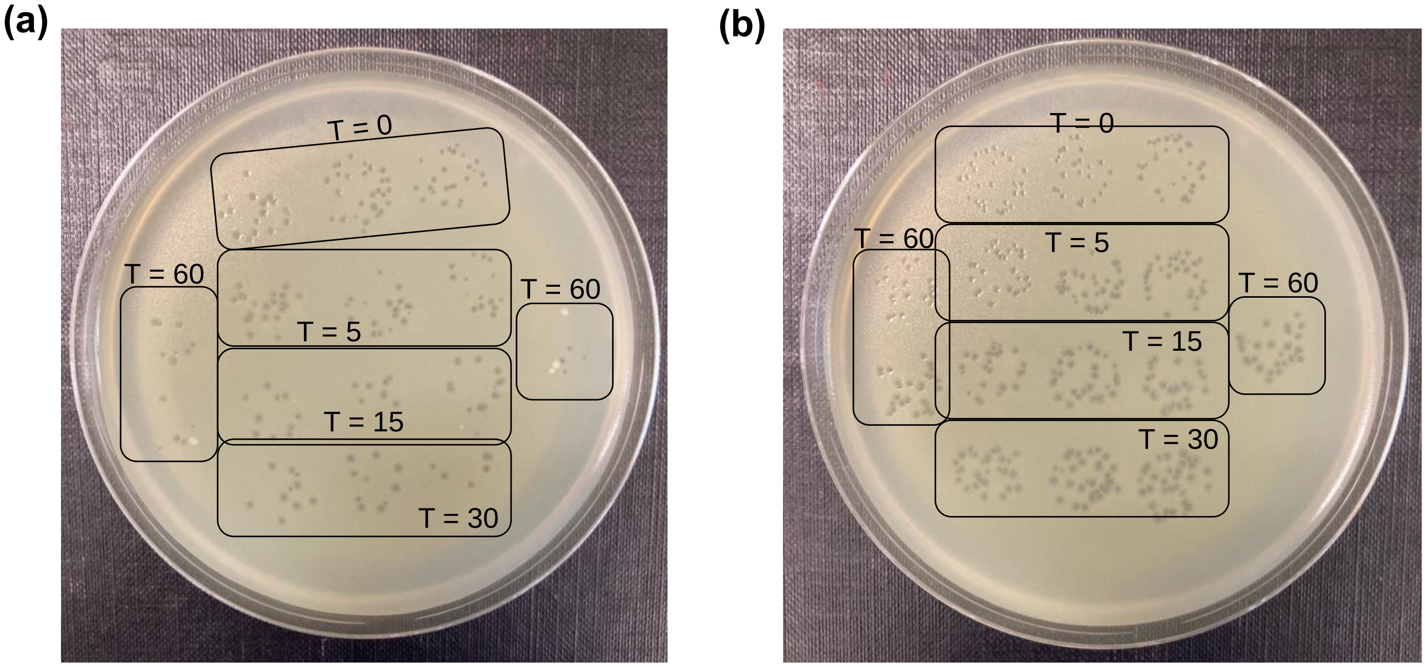


SFig 2. T4 adsorption assay. (a) Plate showing plaques (in triplicates) for phage incubated with overnight *E. coli* culture. (b) Plate showing plaques (in triplicates) for the ‘no bacteria’ control, where phage was added to LB. Time points are shown in minutes.

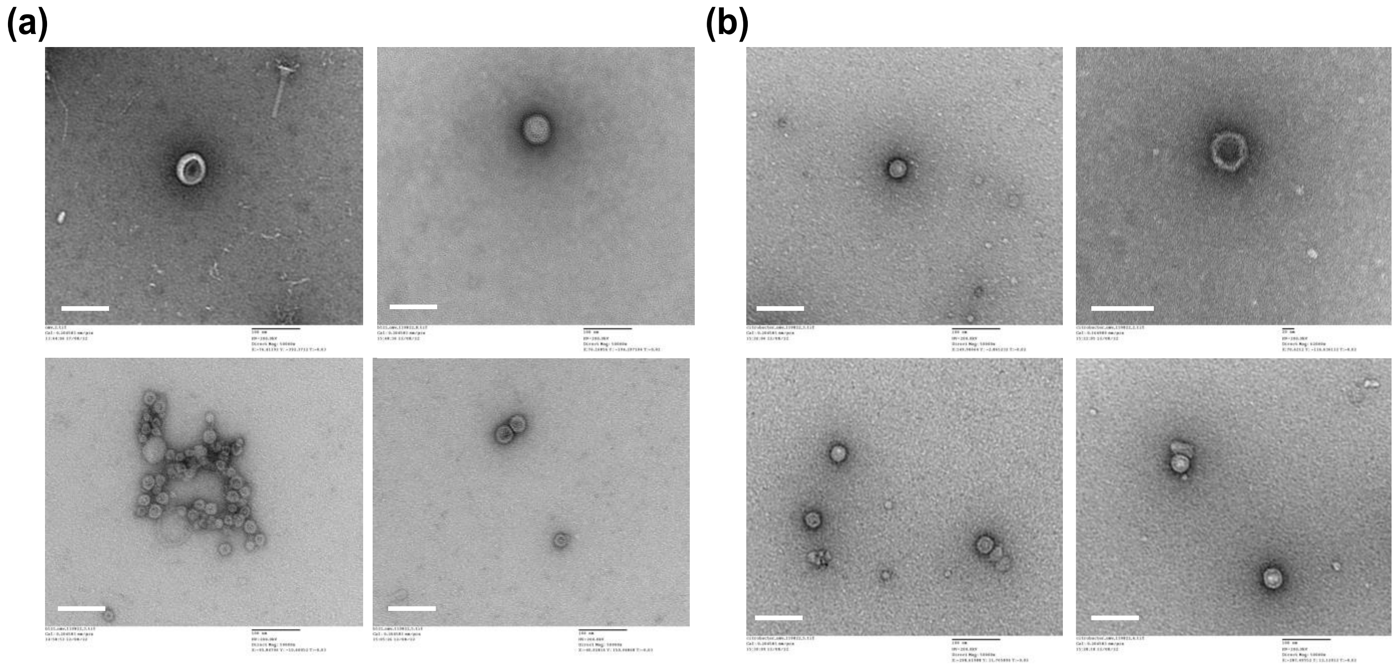

SFig 3. TEM images of OMVs from (a) *E. coli* and (b) *C. rodentium* cells. Scale bars = 100 nm.

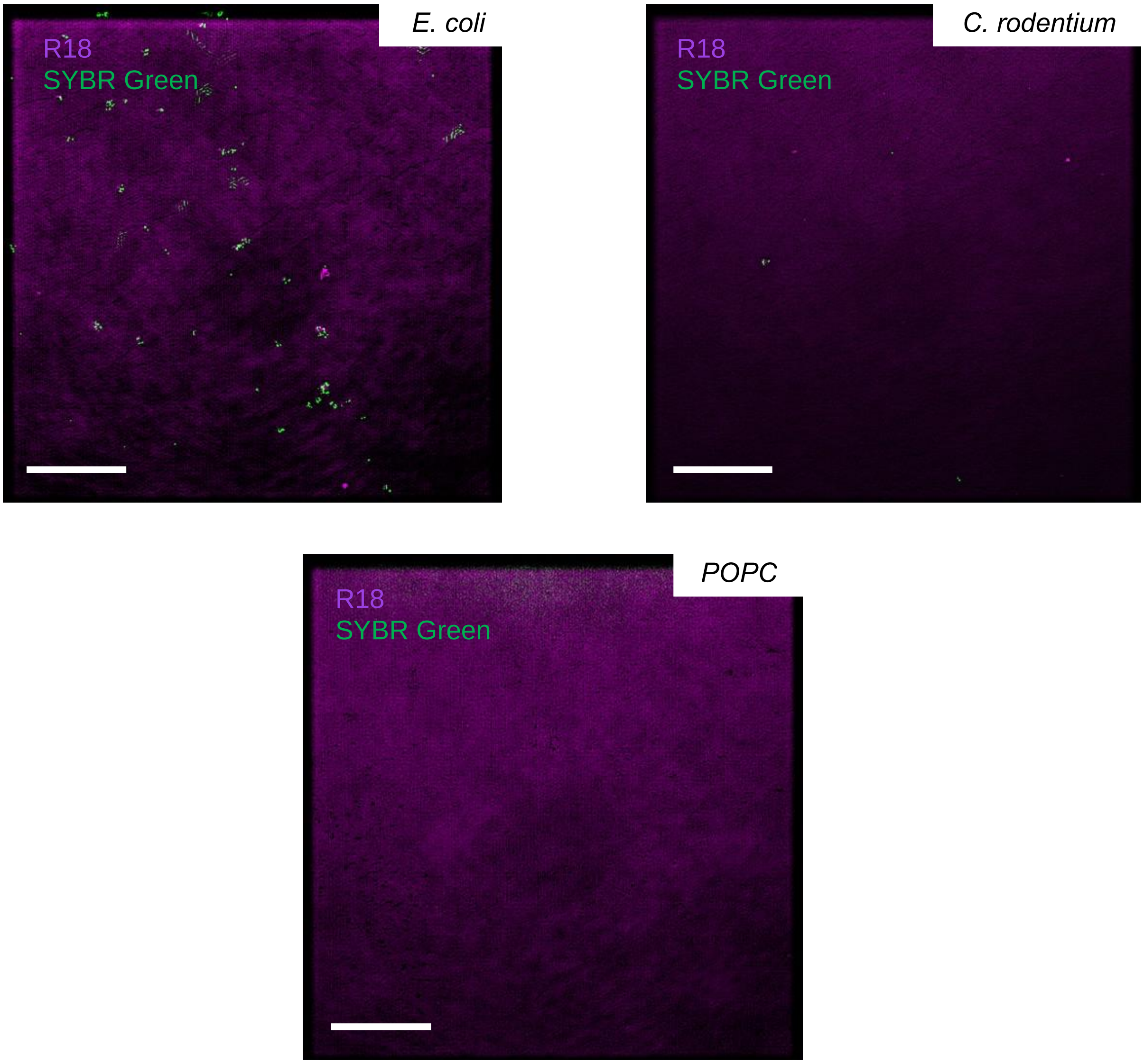


SFig 4. SIM images of the three types of SLBs (*E. coli*, *C. rodentium*, POPC), stained with R18, incubated with SYBR green T4 phage. The areas of green fluorescence in the BL21 SLB are indicative of phage binding. Scale bar = 10 µm.

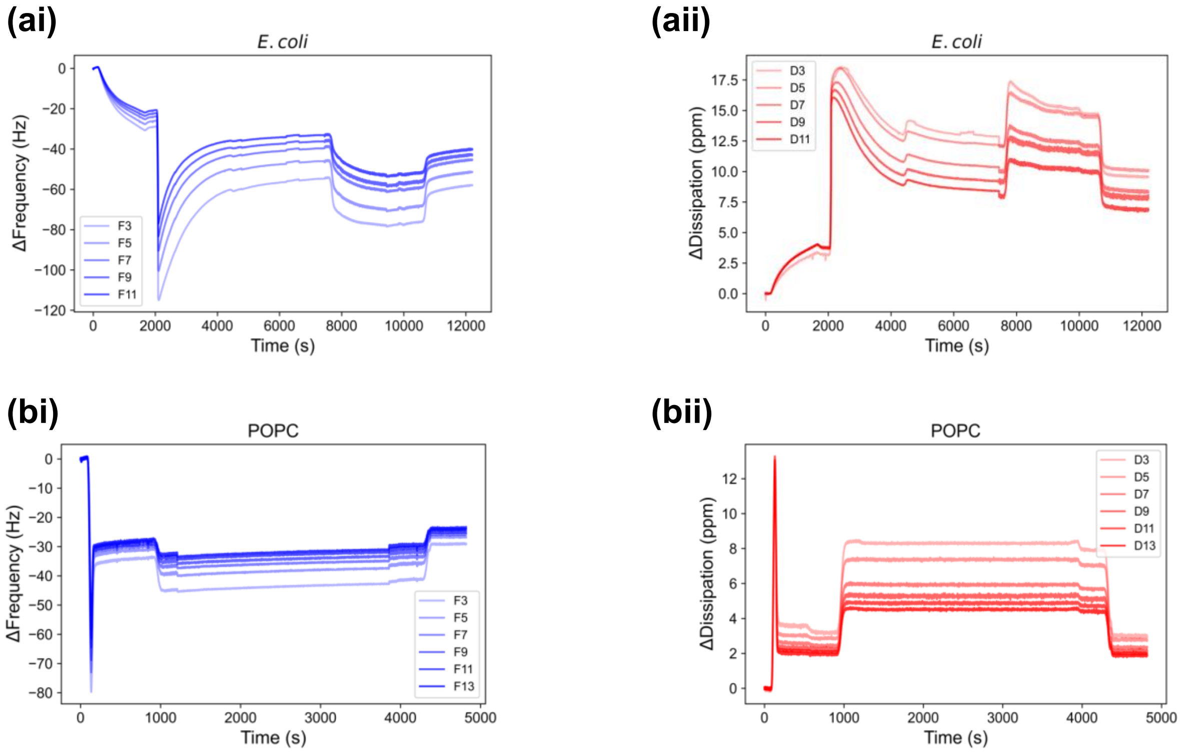


SFig 5. QCM-measurements. (ai) Δf and (aii) ΔD over time for the *E. coli* SLB and phage addition (added at ~ 8000 s, wash step at ~ 10600 s). (bi) Δf and (bii) ΔD over time for the POPC SLB and phage addition (added at ~ 1000 s, wash step at ~ 4300 s).

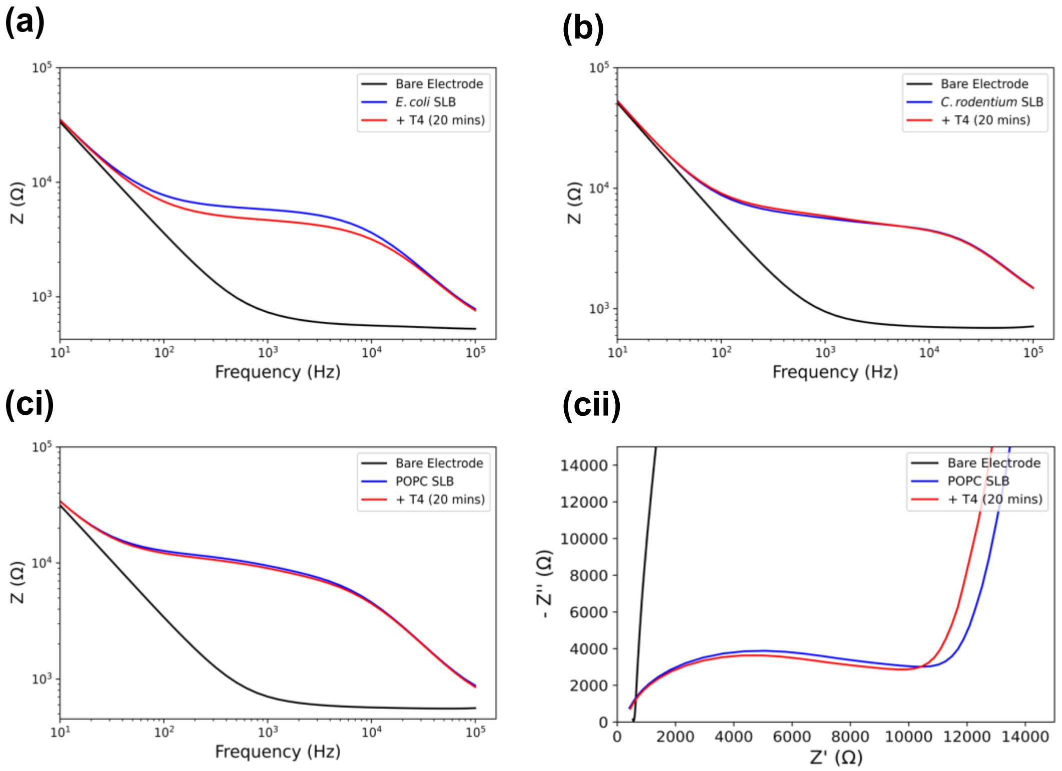


SFig 6. EIS measurements for T4 phage addition to SLBs. Bode plots for (a) *E. coli* and (b) *C. rodentium* SLBs from a representative electrode, corresponding to the Nyquist plots in the main text (figure 4). (ci) Bode and (cii) Nyquist plots from a representative electrode recording for a POPC SLB before and after T4 phage addition.
